# Challenges in Benchmarking Metagenomic Profilers

**DOI:** 10.1101/2020.11.14.382994

**Authors:** Zheng Sun, Shi Huang, Meng Zhang, Qi-Yun Zhu, Niina Haiminen, Anna-Paola Carrieri, Yoshiki Vázquez-Baeza, Laxmi Parida, Ho-Cheol Kim, Rob Knight, Yang-Yu Liu

**Author notes:** These authors contributed equally (Zheng Sun, Shi Huang).

## Abstract

Accurate microbial identification and abundance estimation are crucial for metagenomics analysis. Various methods for classifying metagenomic data and estimating taxonomic profiles, broadly referred to as metagenomic profilers, have been developed. Yet, benchmarking metagenomic profilers remains challenging because some tools are designed to report relative sequence abundance while others report relative taxonomic abundance. Here, we show how misleading conclusions can be drawn by neglecting this distinction between relative abundance types when benchmarking metagenomic profilers. Moreover, we show compelling evidence that interchanging sequence abundance and taxonomic abundance will influence both per-sample summary statistics and cross-sample comparisons. We suggest that the microbiome research community should pay attention to potentially misleading biological conclusions arising from this issue when benchmarking metagenomic profilers, by carefully considering the type of abundance data that was analyzed and interpreted, and clearly stating the strategy used for metagenomic profiling.

---

Identifying microbial species present in complex biological and environmental samples is one of the major challenges in microbiology^1,2^. By directly interrogating the community composition in an unbiased and culture-independent manner, metagenomic sequencing is transforming microbiology by enabling more rapid species detection and discovery^2^. This has a wide range of applications from surveying the bacteria in an environmental soil sample to determining the etiology of an infection from a patient’s blood or stool sample. Such applications drive the development of various computational methods to analyze genomic data generated by metagenomic sequencing to identify all of the species contained in the samples and estimate their relative abundances^2,3^.Those computational methods are broadly referred to as metagenomic profilers.

Following a previous benchmarking study^3^, metagenomic profilers can be categorized based on their reference database type (**Fig.1a**): (1) DNA-to-DNA methods (e.g., Kraken^4,5^, Bracken^6^ and PathSeq^7^), which compare sequence reads with comprehensive metagenome databases; (2) DNA-to-Protein methods (e.g., Kaiju^8^ and Diamond^9^), which compare sequence reads with genomic databases of protein-coding sequences; or (3) DNA-to-Marker methods (e.g., MetaPhlAn^10,11^ and mOTU^12,13^), which only include specific gene families in their reference databases. Note that those metagenomic profilers all rely on reference databases. They should not be confused with *de novo* assembly-based methods that do not use any reference databases^14,15^. Those reference-free binning methods cannot taxonomically classify sequences^14, 15^ and are not directly comparable with the metagenomic profilers evaluated here.

**Figure 1.**
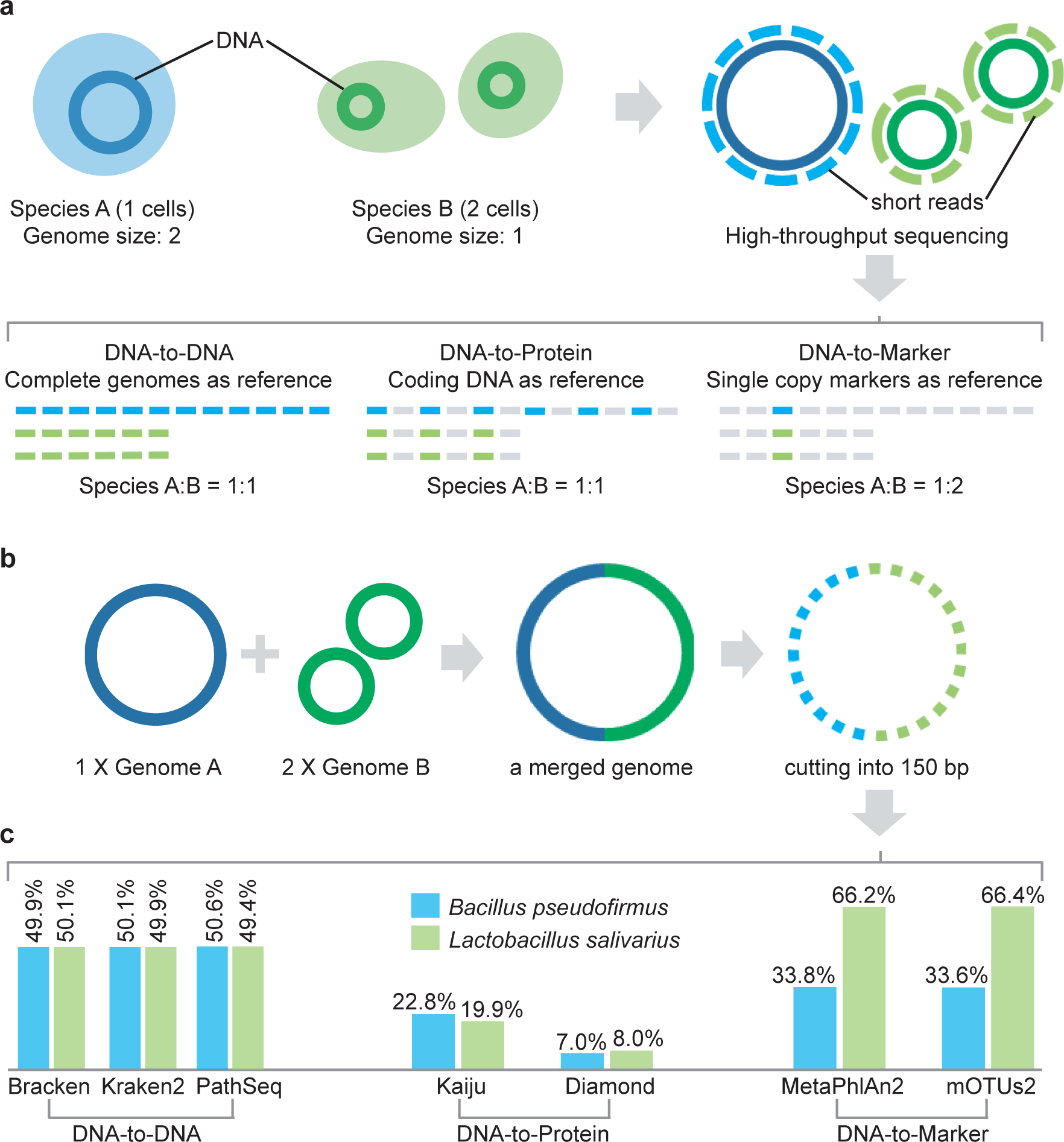
Comparison of profiling results. **a**, Illustration of the reference databases and the default output abundance type for DNA-to-DNA, DNA-to-Protein and DNA-to-Marker profilers on a mixture of two species A (1 cell) and B (2 cells). **b**, A simulated microbial community with only two genomes: *Bacillus pseudofirmus* (genome size 4.2MB) and *Lactobacillus salivarius* (genome size 2.1MB). We merged one copy of *Bacillus pseudofirmus* genome (genome A) with two copies of *Lactobacillus salivarius* genome (genome B) sequences into one metagenome file. Then we sheared the merged metagenomic sequences into 150bp to simulate a typical metagenomic dataset. **c**, Profiling results (default output) of different profilers for the simulated microbial community shown in a. The bar plots show the estimated relative abundance of the two microbial members A and B using different metagenomics profilers.

Many studies have benchmarked metagenomic profilers^3,16-19^, finding that the performance of different profilers varies considerably even on the same benchmark datasets. For example, in a recent benchmarking study^3^, the performance of 20 metagenomic profilers were evaluated based on two key metrics: the area under the precision-recall curve (AUPRC) for organism presence/absence, and the L2 distance between the observed and true relative abundance profiles. It was found that DNA-to-DNA methods were among the best-scoring methods, with typical average L2 distance < 0.1, while DNA-to-Marker methods had much higher L2 distance, indicating less favorable performance.

Here we show that this apparently high performance variation largely arises because the methods report one of two fundamentally different types of relative abundances: *sequence abundance* or *taxonomic abundance*. For example, the raw output of DNA-to-DNA methods is the relative abundance of a given taxon calculated as the proportion of sequences assigned to it out of the total number of sequences, i.e., the sequence abundance. By contrast, DNA-to-Marker methods directly output the relative abundance of each microbial taxon calculated as the number of genomes of that taxon relative to the total number of genomes detected, i.e., the taxonomic abundance. For DNA-to-Protein methods, the output type is the relative sequence abundance of protein-coding sequences^8,9^.

Unfortunately, the distinction between the two types of relative abundances has rarely been carefully considered in previous benchmarking studies. In this paper, we show that the two types of relative abundances are not related by any simple algebraic relation. Moreover, interchanging them leads to very misleading performance assessments of metagenomic profilers. These results imply that many benchmarking results presented in the literature require re-examination. Beyond examining the previous benchmarking results, we further point out the serious issues in microbiome data analysis based on sequence abundances, which are typically produced by DNA-to-DNA methods and have been applied in thousands of published microbiome studies (e.g., Kraken: 1,283 citations; Kraken2: 95 citations; Bracken: 139 citations by November 2020, according to their official websites). We find that microbiome data analysis based on sequence abundance will underestimate (or overestimate) the relative abundances of microbes with smaller (or larger) genome sizes. This will fundamentally affect differential abundance analyses and other analytical methods that rely on accurate counts in their input contingency matrix. Without careful consideration, these issues could impede cross-study comparisons of differentially abundant taxa identified from different methods. We think this point needs more attention from the entire microbiome research community.

## Results

### Illustration of the caveat in benchmarking metagenomic profilers

To illustrate the caveat of confusing sequence abundance and taxonomic abundance in benchmarking metagenomic profilers, we simulated a simple microbial community with only two genomes, where genome A (*Bacillus pseudofirmus*, GCF_000005825.2, size: 4.2MB) is twice the size of genome B (*Lactobacillus salivarius*, GCF_000008925.1, size: 2.1MB), corresponding to **Fig.1b**. In this simulated community, the sequence abundance ratio of genome A: genome B = 1:1, while the taxonomic abundance ratio of genome A: genome B = 1:2. DNA-to-DNA profilers Bracken, Kraken2 and PathSeq reported that this sample contains 49.9% (or 50.1% in Kraken2 and 50.6% in PathSeq) *Bacillus pseudofirmus* and 50.1% (or 49.9% in Kraken2 and 49.4% in PathSeq) *Lactobacillus salivarius*, respectively (**Fig.1c**). DNA-to-Markers profilers MetaPhlAn2 and mOTUs2 reported the relative abundance of *Bacillus pseudofirmus* as 33.8% (or 33.6%) and *Lactobacillus salivarius* as 66.2% (or 66.4%, **Fig.1c**), respectively. This simple example clearly illustrates how the sequence abundance profile produced by DNA-to-DNA profilers does not represent the true taxonomic abundance of a microbiome sample.

Note that for this simple synthetic community, DNA-to-Protein profilers Kaiju and Diamond reported the relative abundance of *Bacillus pseudofirmus* as 22.8% (or 7.0%) and *Lactobacillus salivarius* as 19.9% (or 8.0%), respectively (**Fig.1c**). Besides the false positives (57.3% in Kaiju and 85.0% in Diamond), the ratio between the relative abundances of the two species is roughly 1:1, indicating the methods are indeed reporting sequence abundance. However, these classifiers reported a large number of false positive species identified due to the conservation of protein sequence^20^. Going forward, we will focus on benchmarking the DNA-to-DNA and DNA-to-Markers methods.

### No simple algebraic relation between the two types of relative abundances

We emphasize that mathematically there is no simple algebraic relation between the two types of relative abundances, even in the ideal case (when all genomes/taxa are known). Denote *R_i_* as the number of metagenomic reads assigned to the genome of a microbial taxon *i* with genome size *L_i_* and ploidy *P_i_* (i.e., the number of copies of the genome in one cell, however most methods did not consider the ploidy into the abundance estimation as the information is still lacking for many genomes). The number of microbial cells classified as taxon *i* is then given by *C_i_ = R_i_*/(*L_i_P_i_*). Let *n* be the number of identified taxa in the sample. Then the sequence abundance of taxon *i* is given by

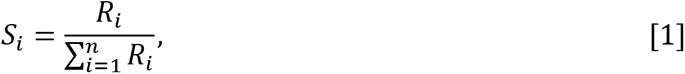

and its taxonomic abundance is given by

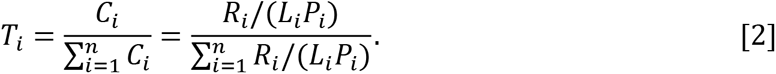

Eqs.[1-2] imply that as long as *L_i_* and *P_i_* vary across different taxa in a community, *S_i_* and *T_i_* are not connected by any simple algebraic relation.

The variation of genome size *L_i_* of different taxa can be very large. Indeed, in the recently updated microbial genome database (NCBI RefSeq, 2020 Nov 6^th^), the sizes of fully sequenced and assembled microbial genomes vary considerably (**Fig.2a**). For example, just within the bacteria kingdom, the genome size variation can be more than 100-fold, e.g., *Candidatus Nasuia deltocephalinicola* (GCF_000442605.1) with 112,091 bp vs. *Sorangium cellulosum* (GCF_000418325.1) with 14,782,125 bp. Therefore, microbial genome sizes could vary radically within a single microbiome sample, including when viruses (which tend to have shorter genomes, Fig. 2a) are analyzed together with bacteria in shotgun metagenomics.

**Figure 2.**
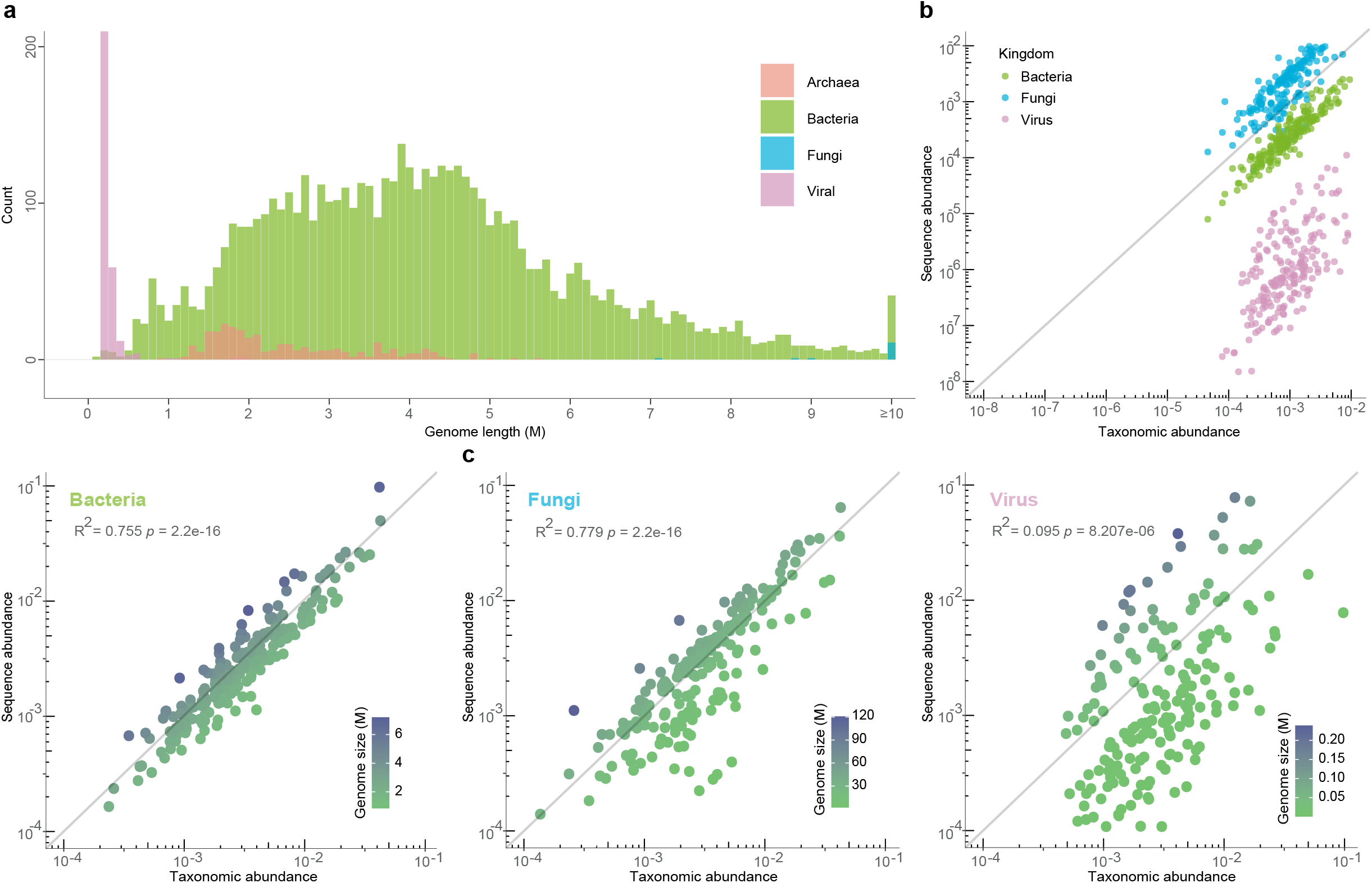
Correlation between sequence abundance and taxonomic abundance in synthetic profiles based on different kingdoms. **a**, Genome size distribution of microorganisms calculated from the microbial genome database (NCBI RefSeq 2020 Nov 6^th^) that includes 171,927 bacteria, 293 fungi, 945 archaea, and 9,362 viruses. **b**, The scatter plot shows the correlation between taxonomic abundance (x axis) and sequence abundance (y axis) of 600 randomly selected species in a simulated profile which includes bacteria (species number=200), fungi (species number=200) and virus (species number=200). **c,** Each scatter plot shows the correlation between taxonomic abundance (x axis) and sequence abundance (y axis) of 200 randomly selected species in three simulated profiles which represent different kingdoms e.g. bacteria, fungi, and virus.

Regrading ploidy *P_i_*, although prokaryotes are usually thought to contain one copy of a circular chromosome, previous studies have demonstrated that many species of archaea and bacteria are polyploid and can contain more than ten copies of their chromosome^21^. In fact, extreme polyploidy has been observed in a large bacterium *Epulopiscium*, which contains tens of thousands of copies of its genome^22^.

The variations in *L_i_* and *P_i_* drive the theoretical distinction between sequence abundance and taxonomic abundance. This point can be seen clearly from simulated microbial communities based on the NCBI RefSeq database. As shown in **Fig.2b**, where we investigate a complex microbial community consisting of all different kingdoms of microbes (fungi, bacteria and virus), *S_i_* tends to overestimate the abundances of species with larger genome sizes (e.g., fungi) and underestimate the abundances of species with smaller genome sizes (e.g., viruses). This is true even if we investigate a community consisting of microbes from the same kingdom (**Fig.2c**). Note that here, for the sake of simplicity, in our simulations we did not consider the variation of ploidy, but only focused on the variation of genome sizes. Hence, the demonstrated difference between sequence abundance and taxonomic abundance is conservative.

In reality, unknown genomes/taxa will further complicate the relation between Si and Pi, and affect metagenomic profiler benchmarking on real data (because different profilers handle unknown genomes/taxa differently). Moreover, instead of converting *S_i_* to *T_i_* through *L_i_* and *P_i_* correction, DNA-to-Marker methods directly calculate *T_i_* as the ratio of sequence coverage of single-copy marker genes of each taxon to that of all taxa. This also affects the metagenomic profiler benchmarking.

### Benchmarking results depend on the abundance type

To further illustrate the problem of mixing sequence abundance and taxonomic abundance in benchmarking metagenomic profilers, we simulated metagenomic sequencing reads for 25 communities from distinct habitats (e.g., gut, oral, skin, vagina and building, five communities for each habitat, see **Methods**). To avoid database biases of different metagenomic profilers, the selection of genomes for simulated data was based on the intersection between MetaPhlAn2, mOTUs2 reference database, and Kraken2 reference database (which was also used by Bracken). Then we calculated the dissimilarity or distance between the ground truth abundance profiles and the estimated ones from different profilers, based on the following five measures: Bray-Curtis dissimilarity (BC), L1 distance, L2 distance, root Jensen-Shannon divergence (rJSD), and robust Aitchison distance (rAD)^23^ (**Fig.3a,b**). Note that the Aitchison distance (based on centered log-ratio transform) is a compositionally aware distance measure^23^. However, it suffers from the inflated zero counts in microbiome data because log-transform of zero counts is undefined unless arbitrary pseudocounts are added to each taxon. Here the calculation of rAD does not involve any pseudocounts, and it naturally avoids the issue of dealing with sparse zero counts using the classical Aitchison distance^23^.

**Figure 3.**
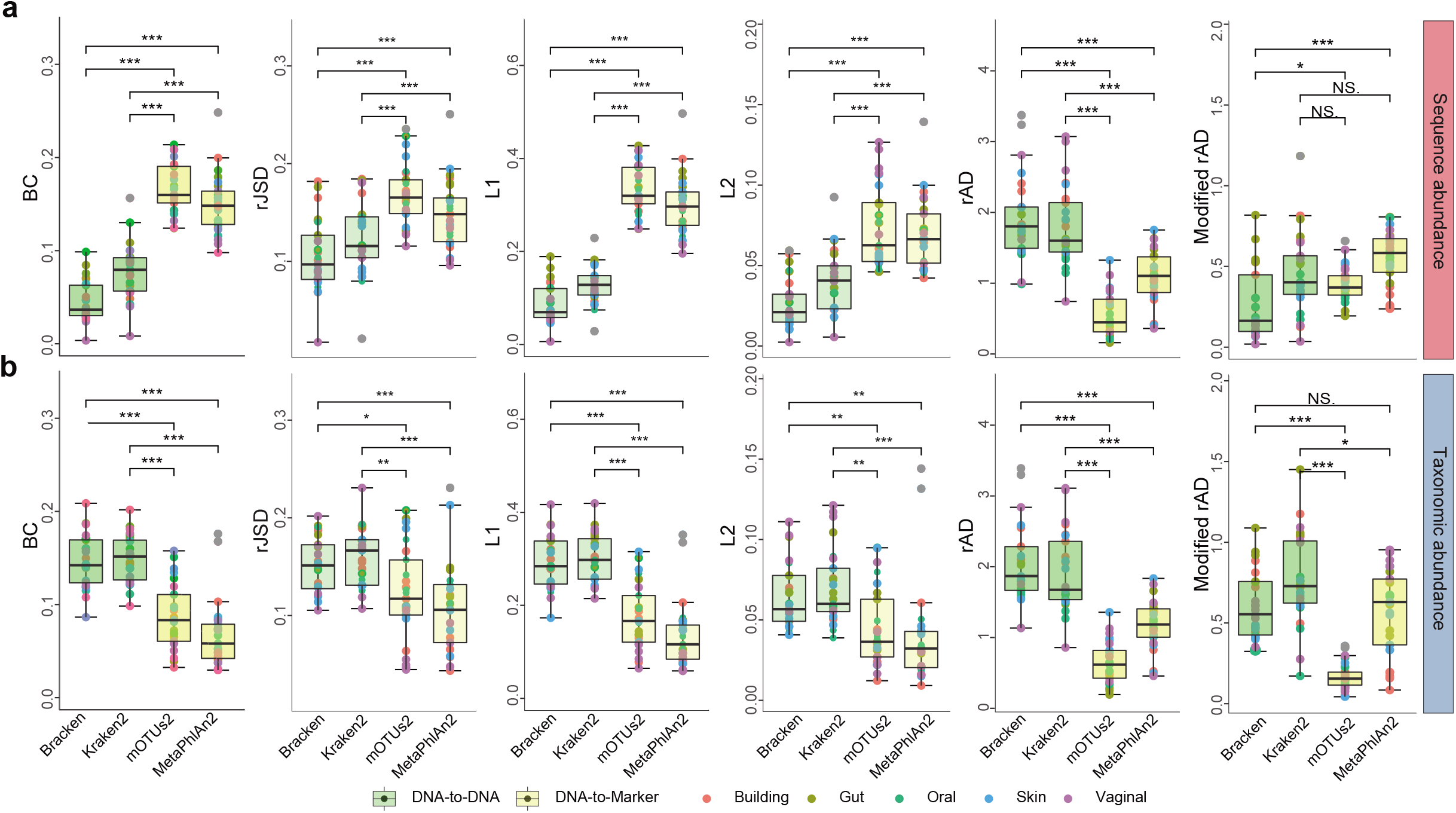
Differential benchmarking results of four representative metagenomics profilers using two types of relative abundance as ground truth: **a**, sequence abundance and **b**, taxonomic abundance. The boxplots indicate the dissimilarities based on L1, L2, root Jensen-Shannon divergence (rJSD), Bray-Curtis (BC), and robust Aitchison distance (rAD) between the ground-truth profiles and the profiles predicted by different metagenomics profilers (Bracken, Kraken2, mOTUs2, and MetaPhlAn2) at the species level. For each metagenomic profiler, we performed the dissimilarity calculations based on 25 simulated microbial communities from five representative environmental habitats (gut, oral, skin, vagina and building) separately. Note that for each profiler based on any evaluation metric, its performance variation across different synthetic communities is due to microbiome complexity difference (e.g. species composition and richness). The asterisks in the boxplots refer to the statistical significance: “*” refers to p-value <0.05, “**” refers to <0.01, “***” refers to < 0.001.

We found that for BC, L1, L2 and rJSD, if the sequence abundance is used as the ground truth, Bracken and Kraken2 outperform MetaPhlAn2 and mOTUs2; while if the taxonomic abundance is used as the ground truth, MetaPhlAn2 and mOTUs2 outperform Bracken and Kraken2. Interestingly, with rAD as the evaluation metric, regardless of if sequence or taxonomic abundance profiles were taken as the ground truth, mOTUs2 and MetaPhlan2 always outperform Bracken and Kraken. This could be due to the compositionally aware distance measure rAD weighing low-abundance taxa more than the other measures. To test this idea, we sought to rule out the bias introduced by false positives and calculated rAD based on taxonomic profilers where false positives have been removed (**Methods**). This is denoted as modified rAD in **Fig.3**. We found that, with the modified rAD as the evaluation metric, the benchmarking result is the same as that of using BC, L1, L2 and rJSD, or their modified versions (**Fig.S1**). We always found the same pattern: if the sequence abundance is used as the ground truth, Bracken and Kraken2 outperform MetaPhlAn2 and mOTUs2; while if the taxonomic abundance is used as the ground truth, MetaPhlAn2 and mOTUs2 outperform Bracken and Kraken2. This result strongly indicates that the benchmarking result of metagenomic profilers depends on the selected abundance type.

We emphasize that the above contradicting performance evaluations due to different abundance types cannot be detected by using the AUPRC metric, because the calculation of the Precision-Recall Curve only concerns the difference of presence/absence patterns in the ground truth and predicted abundance profiles. By definition, the ground truth sequence abundance and taxonomic abundance profiles of our simulated microbiome samples share exactly the same presence/absence pattern.

Moreover, we emphasize that even though the five distance/dissimilarity measures (BC, L1, L2, rJSD, and rAD) all showed the similar results in the performance evaluation (after the removal of false positives), L2 was not designed for compositional data analysis. To investigate whether the discriminating power of these distance measures for the two sequence types persists with varied microbial diversity, we simulated a set of abundance tables (for both taxonomic abundance and sequence abundance) with different species counts ranging from 10 to 500 (see **Methods**). We then calculated the distance or dissimilarity between the sequence abundance and taxonomic abundance profiles (**Fig.4**). We found that with an increasing number of species, L2 keeps decreasing while L1, BC, rJSD and rAD can still distinguish the two abundance types. This result suggests that L2 distance cannot discriminate the two types of relative abundances in microbiome samples of high species richness. This might be due to the fact that L2 distance is not appropriate for compositional data analysis.

**Figure 4.**
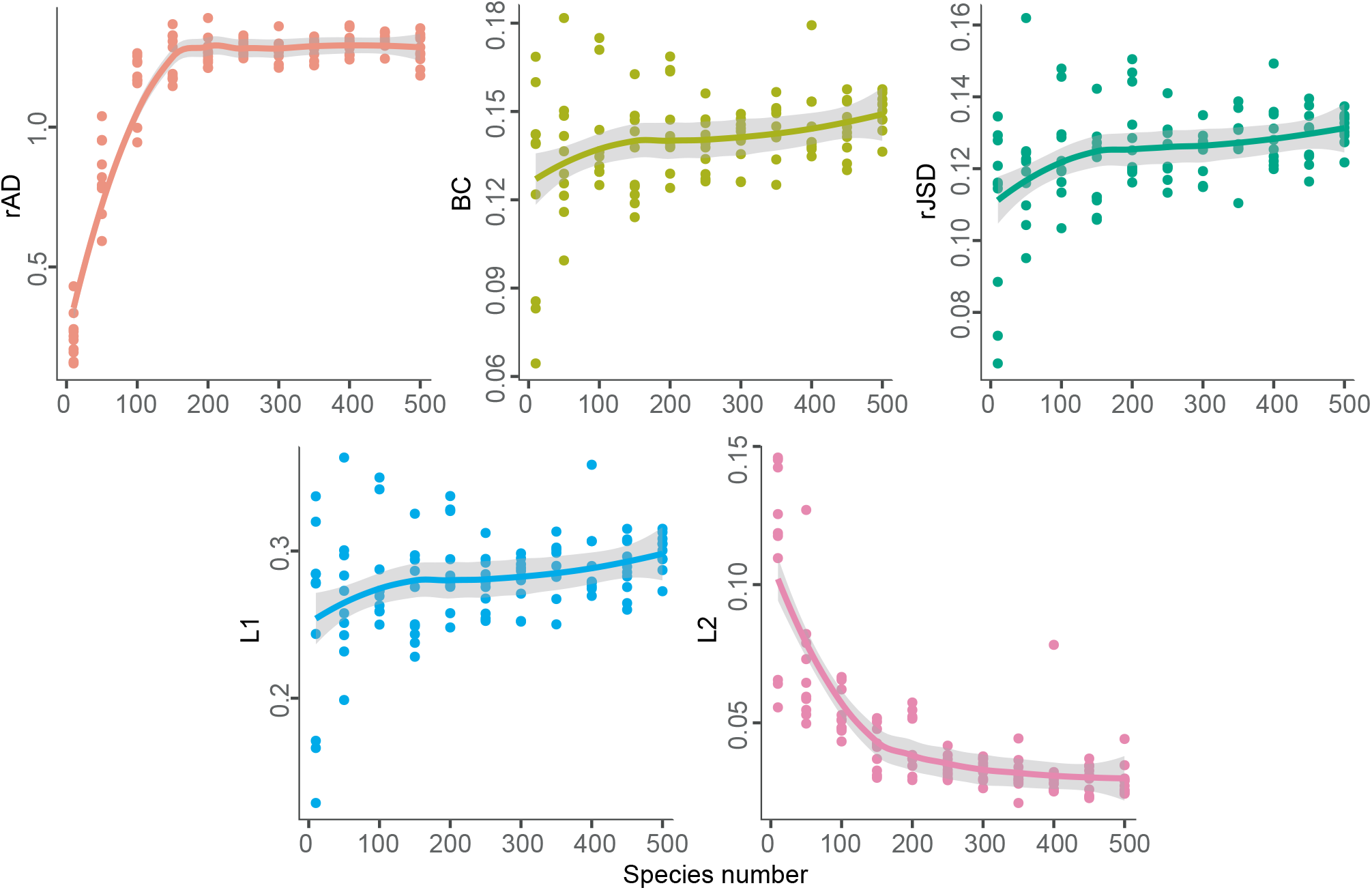
Dissimilarity between sequence abundance and taxonomic abundance with varied species number measured by different distance measures. For each species number, we simulated 10 repeats of profiles. The distance/dissimilarity was then measured by different measures: rAD (red), L1 (blue), L2 (purple), Bray-Curtis (yellow) and rJSD (green). rAD between these types of abundance profiles positively correlated with the species richness when < 200 microbial species presented in a community, yet saturated after the number of species reaching 200. L1, BC and rJSD can also reveal the difference between the two abundance types yet they were not affected by the species-level richness. L2 distance between the two abundance types dramatically dropped with the increase in the species-level richness. In the complex community with the number of species over 200, L2 distance metric almost lost the discriminatory power of these two abundance profiles.

### Impact of abundance type on the alpha diversity calculation

Interchanging sequence abundance and taxonomic abundance strongly influences per-sample summary statistics. To demonstrate this issue, we simulated 500 abundance profiles representing microbiota from distinct habitats (gut, oral, skin, vagina, and building, 100 profiles for each, see **Methods**) with known sequence abundance and taxonomic abundance profiles. We found that the Shannon and Simpson indices calculated from taxonomic abundances are significantly higher than those calculated from sequence abundances (*p*<0.001, Wilcoxon rank-sum test) regardless of the habitat (**Fig.5**). Moreover, when ranking the samples according to their alpha diversity measures calculated from sequence abundance and from taxonomic abundance, the orderings are not fully concordant with each other (Spearman correlation of the rank vectors is 0.929 ± 0.020 for Shannon index and 0.835±0.042 for Simpson index). These results suggest that alpha diversity calculations and comparisons can be strongly affected by the type of relative abundance used.

**Figure 5.**
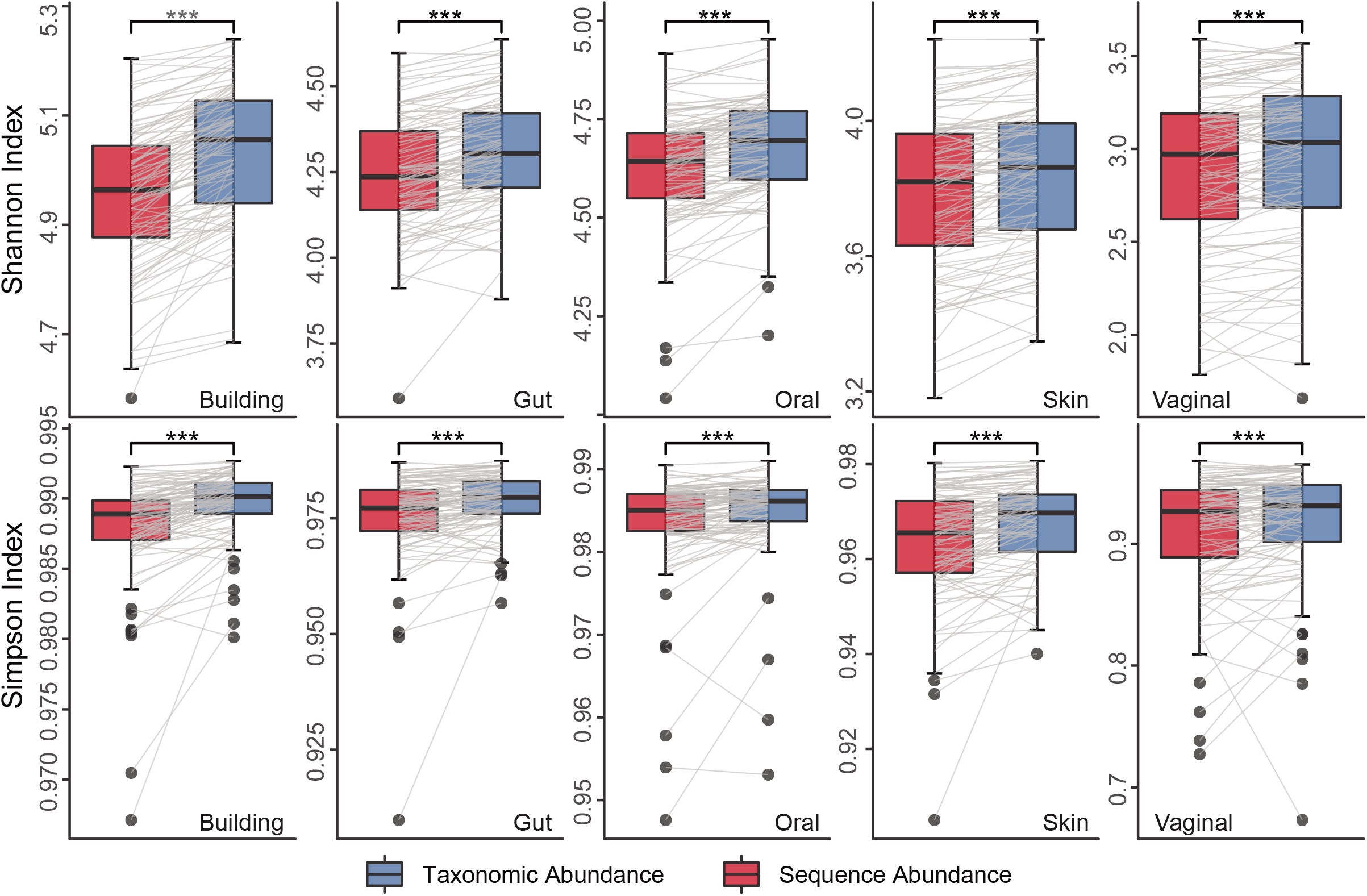
Alpha diversity based on sequence abundance and taxonomic abundance. Alpha diversity (Shannon index and Simpson index) based on ground truth of simulated data from different habitats revealed the influence of abundance types. The index within sample between two abundance type were connected to illustrate the change trend of the indices, the asterisks representing significantly differences are based on paired Wilcoxon test, “***” refers to P < 0.001.

### Impact of abundance types on the beta diversity and ordination analyses

To check if mixing sequence abundance and taxonomic abundance will also influence between-sample attributes such as beta diversity and ordination analyses, we reanalyzed the 500 samples generated for Fig.5. In order to quantify the influence on beta diversity introduced by abundance type, Mantel test^24, 25^ was employed to compare the beta-diversity (in terms of BC, rJSD, L1, L2 and rAD) calculated from the taxonomic abundance and sequence abundance profiles of those samples (see **Methods**). Interestingly, regardless of the species richness in the habitats, the abundance type has some influence on the cross-sample comparisons based on the BC, rJSD and L1 measures (Spearman coefficient r=0.944±0.006, 0.947±0.009, 0.944±0.006, respectively; p-value =1e-4 for all), but affects the L2 and rAD measures more strongly (r =0.844±0.026, 0.519±0.137, respectively; p-value=1e-4 for both). Moreover, we found that species richness of samples associates with the correlation coefficient in the rAD calculation. These results demonstrate the inconsistent relative relationships between samples that are introduced by different abundance types in beta diversity calculation.

We then performed ordination analyses using four different methods: Non-metric Multidimensional Scaling (NMDS)^26^, Principal Coordinates Analysis (PCoA)^27^, t-distributed stochastic neighbor embedding (t-SNE)^28^, and Uniform Manifold Approximation and Projection (UMAP)^29^. We found that, regardless of the distance/dissimilarity measures used (e.g. rJSD, BC and rAD), taxonomic abundance and sequence abundance profiles are drastically different in all the four ordination results (**Fig.6, Figs.S2-S3**). Procrustes analysis was then employed to analyze the congruence of two-dimensional shapes produced from superimposition of ordination analyses from two datasets^30, 31^. Indeed, Procrustes analysis revealed very low similarity between the ordination results calculated from sequence and taxonomic abundance (Fig.6, Figs.S2-S3, Monte Carlo p-value<0.05). These results indicate that both beta diversity (especially for L2 and rAD) and ordination analyses can be heavily affected by the relative abundance type used.

**Figure 6.**
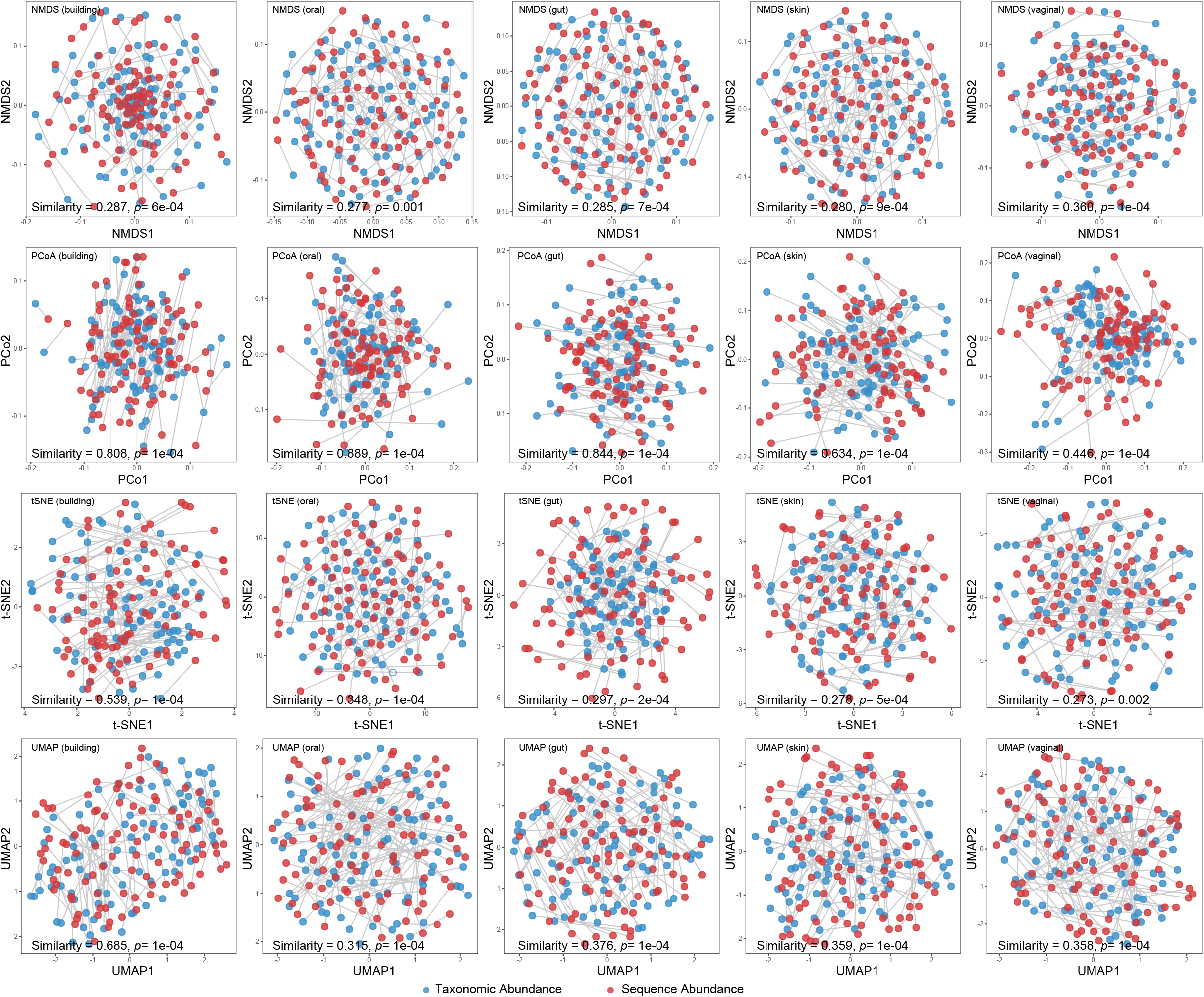
Ordination analyses of simulated profiles based on rJSD. Scatter plots of NMDS, PCoA, t-SNE and UMAP illustrate the dissimilarities between the sequence abundance (red dots) and taxonomic abundance (blue dots), which are the ground truth of the simulated 100 gut profiles. Root Jensen-Shannon divergence (rJSD) was used to for the ordination analyses. The plots of the ordination analyses based on sequence abundance and taxonomic abundance were adjusted to overlap with each other first, then the similarity was calculated by the Monte-Carlo test. The two abundance types from the same profile were connected using grey lines to show the change of its position.

## Discussion

Taken together, these analyses emphasize the importance of differentiating between relative sequence abundance and relative taxonomic abundance in metagenomic profiling. Ignoring this distinction can potentially underestimate the relative abundance of organisms with small genome sizes. Sequence abundances are typically produced by DNA-to-DNA or DNA-to-Protein methods, which rely on microbial genomes or genes as the reference database, report relative sequence abundance, i.e. the fraction of sequence reads assigned to each entity in the database. By contrast, DNA-to-Marker methods output relative taxonomic abundance representing the fraction of each detected taxon.

Our results demonstrate that misleading performance assessment of metagenomic profilers and spurious alpha and beta diversity patterns can arise from interchanging sequence abundance with taxonomic abundance. For alpha diversity, Shannon index and Simpson index are not simply higher based on taxonomic abundance than that based on sequence abundance, the relative ranking of alpha diversity is not consisting in the two abundance types either. Dramatic changes in the relative position between samples are also shown in the ordination analysis. Therefore, interchanging abundance types could have a deleterious effect on the interpretation of alpha and beta diversity analyses and meta-analyses.

The distinction between the two types of relative abundances was known to the field of microbiome research (at least to the developers of various metagenomic profilers), and has been conceptually considered in some benchmark studies (e.g., CAMI^19^). However, the consequences of ignoring this distinction for benchmarking metagenomic classifiers and per-sample summary statistics have not been quantitatively studied or clearly illustrated so far. In particular, the vast majority of users of those metagenomic profilers should be clearly aware of the distinction between sequence abundance and taxonomic abundance, and of the consequences of ignoring this distinction in selecting metagenomics tools, data interpretation, and cross-study comparison of differentially abundant taxa identified by different tools.

In summary, we suggest that the microbiome research community should pay more attention to potentially misleading biological conclusions arising from this issue by carefully considering which type of abundance data was analyzed and interpreted, and, going forward, the strategy used for taxonomy assignment should be clearly represented. We also suggest that, in future development or evaluation of metagenomic profilers, both types of relative abundance should be strictly distinguished, and both should be reported. This would substantially improve the comparison of abundance estimations of metagenomic profilers and enhance the reproducibility and biological interpretation of microbiome studies.

## Methods

### Simulation of microbiome profiles

In the simulation of microbiome profiles based on different species counts (from 10 to 500), the abundance was created randomly from a log-normal distribution using “rlnorm” function in R language with parameters: meanlog = 0 and sdlog = 1, and 10 repeats were simulated for each species count. In the simulation of microbiome profiles for alpha diversity calculation, 100 profiles were simulated for each habitat, and species counts in different habitats were set up as: 10-50 (vaginal), 50-100 (skin), 100-150 (gut), 150-200 (oral), 200-300 (building). The representative species in each specific habitat were selected based on the set of microbial species identified in the HMP^32^ and by Hsu et al.^33^.

### Simulation of sequencing reads

Firstly, the 25 microbiome profiles (five for each habitat) were simulated using the above method. Then the simulation of sequencing data is illustrated as the process in Fig.1a: Given a specified species composition (taxonomic abundance), their sequence abundance can be inferred accordingly (taxonomic abundance equals to sequence abundance divide by their genome length) and “Wgsim” (https://github.com/lh3/wgsim) was then used (with default parameters) to simulate the sequences. The selection of genomes for simulated data was based on the intersection between MetaPhlAn2 and mOTUs2 reference database and Bracken’s database to avoid database biases.

Currently, there are many more DNA-to-DNA profilers (e.g., Bracken and Kraken2) than DNA-to-Marker profilers (e.g., MetaPhlAn2 and mOTU2). In this paper we focused on two DNA-to-DNA profilers for the following reasons. First, as representative DNA-to-DNA methods, Bracken and Kraken/Kraken2 demonstrated the best performance in previous benchmarking studies^4, 6, 34^, and have been cited in more than one thousand microbiome studies. Second, mOTU2 and MetaPhlAn2 do not support custom reference databases, and the reference database is a critical factor affecting profiler performance. As such we decided to use the intersection of organisms in mOTU2, MetaPhlaAn2, and Kraken2 reference databases as the source for our simulation data. Introducing more DNA-to-DNA profilers could further reduce the reference database size of the simulated data and affect the diversity of genome sizes (**Fig.S4**).

### Alpha and beta diversity calculation

Alpha diversity calculation e.g. Shannon and Simpson indices were performed in R language by the “Vegan 2.5-6” package. As for the beta diversity, we employed “Vegan 2.5-6” for distance/dissimilarity calculation e.g. L1 (“Manhattan” in vegdist function), L2 (“Euclidean”) and BC (“Bray”), while rJSD and rAD were calculated by self-programmed script (see **code availability**). In the ordination analyses, R packages “ade4 1.7-15”, “Rtsne 0.15”, “ape 5.4-1” and “umap 0.2.6.0” were used to conduct the NMDS, t-SNE, PCoA and UMAP analyses separately. Since the iterative algorithm of NMDS, t-SNE and UMAP find different solutions depending on the starting point of the calculation (which is a randomly chosen configuration) we performed 101 repeats of NMDS, t-SNE, UMAP and their Procrustes test, the median result (sorting by the Mote-Caro test) was selected for presentation of similarity and p-value in **Fig.6, Fig.S2** and **Fig.S3**. The ordination analyses based on the ground truth of the sequence abundance and taxonomic abundance for the 500 profiles (from five habitats) were conducted separately before Procrustes analysis.

### Robust Aitchison distance calculation

We applied DEICODE (https://github.com/biocore/DEICODE) to calculate the robust Aitchison distance (rAD) to benchmark the performance of metagenomics profilers. DEICODE represents a form of Aitchison Distance that is robust to high levels of sparsity. It preprocesses the compositional data using the centered log-ratio (CLR) transform only on the non-zero values of the data (hence no pseudo counts are used). Then it performs dimensionality reduction through robust PCA based on the non-zero values of the data. The Euclidean distance of the robust CLR-transformed abundance profiles (i.e., rAD) was finally employed to evaluate the performance of metagenomic profilers. To avoid the impact of false positives on the benchmarking results, we further filtered out false positives in all output taxonomic profiles (Kraken2: 29.26%±12.13%; Bracken: 36.91%±12.11%; mOTUs2: 11.47%±4.62%; MPA2: II. 29%±4.19%) and compared the performance of different profilers using rAD calculated from the true positives only. This is termed as the modified rAD in **Fig.3**. For other evaluation measures, the same procedure was performed and presented in **Fig.S1**.

### Mantel Test

Mantel test was used as a correlation test to determine the correlation between two beta diversity (BC, rJSD, L1, L2 and rAD) matrices based on sequence abundance and taxonomic abundance. In order to calculate the correlation, the matrix values of both matrices are ‘unfolded’ into long column vectors, which are then used to determine correlation. Permutations (n=9999) of one matrix are used to determine significance. Whether distances between samples in one matrix are correlated with the distances between samples in the other matrix is revealed by the p-value.

### Procrustes analysis

Procrustes analysis (by R package “ade4 1.7-15”) typically takes as input two coordinate matrices with matched sample points, and transforms the second coordinate set by rotating, scaling, and translating it to maximize the similarity between corresponding sample points in the two shapes. It allows us to determine whether we would come to same conclusions on the beta diversity, regardless of which distance/dissimilarity measure was used to compare the samples. To assess the significance level of observed similarity between two matrices, empirical p-values are calculated using a Monte Carlo simulation. Basically, sample labels are shuffled in one of the coordinate matrices, and then the similarity between them is re-computed for 9999 times. Here, similarity is calculated as the sum of the squared residual deviations between sample points for each measurement. The proportion of similarity values that are equal to or lower than the observed similarity value is then the Monte Carlo or empirical p-value.

## Data availability

All the simulated datasets can be downloaded here: https://figshare.com/projects/Challenges_in_Benchmarking_Metagenomic_Profilers/79916.

## Code availability

R scripts used in this paper is available at https://github.com/shihuang047/re-benchmarking.

## Acknowledgements

Research reported in this publication was supported by grants R01AI141529, R01HD093761, UH3OD023268, U19AI095219, and U01HL089856 from National Institutes of Health. This work was also supported by IBM Research through the AI Horizons Network, UC San Diego AI for Healthy Living program in partnership with the UC San Diego Center for Microbiome Innovation.

## Author contributions

Y.-Y.L. and R.K. conceived and designed the analysis. Z.S. and H.S. led the analysis. M.Z., Q.Z., N.H., A.-P.C., Y.V., L.P., and H.-C.K. contributed evaluation strategies. All authors analyzed the results. Z.S., H.S., Y.-Y.L., and R.K. wrote the paper. All authors edited the paper.

