## Supplemental figures for "Challenges in Benchmarking Metagenomic Profilers"

**Figure S1.** Differential benchmarking results based on modified distance measures using two types of relative abundance as ground truth.

**Figure S2.** Ordination analyses of simulated profiles based on BC.

**Figure S3.** Ordination analyses of simulated profiles based on rAD.

**Figure S4.** Genome size distribution of bacteria in the reference databases used in Kraken2, MetaPhlAn2, and mOTUs2.

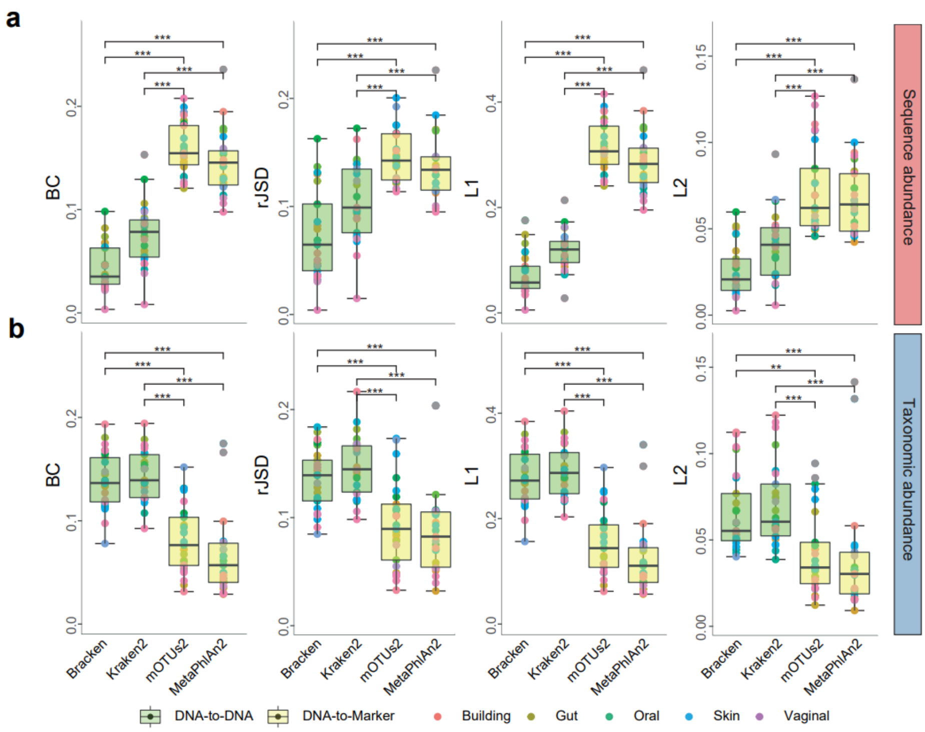


**Figure S1. Differential benchmarking results based on modified distance measures using two types of relative abundance as ground truth**: **a**, sequence abundance and **b**, taxonomic abundance. The boxplots indicate the dissimilarities (based on modified L1, modified L2, modified root Jensen-Shannon divergence (modified rJSD) and modified Bray-Curtis (modified BC) between the ground-truth profiles and the profiles predicted by different metagenomics profilers (Bracken, Kraken2, mOTUs2, and MetaPhlAn2) at the species level. The modified distances were calculated based on the true positives in the resulting profiles as compared to the ground truth. For each metagenomic profiler, we performed the dissimilarity calculations based on 25 simulated microbial communities from five representative environmental habitats (gut, oral, skin, vagina and building) separately. The asterisks in the boxplots refer to the statistical significance: "*" refers to p-value <0.05, "**" refers to <0.01, "***" refers to < 0.001.

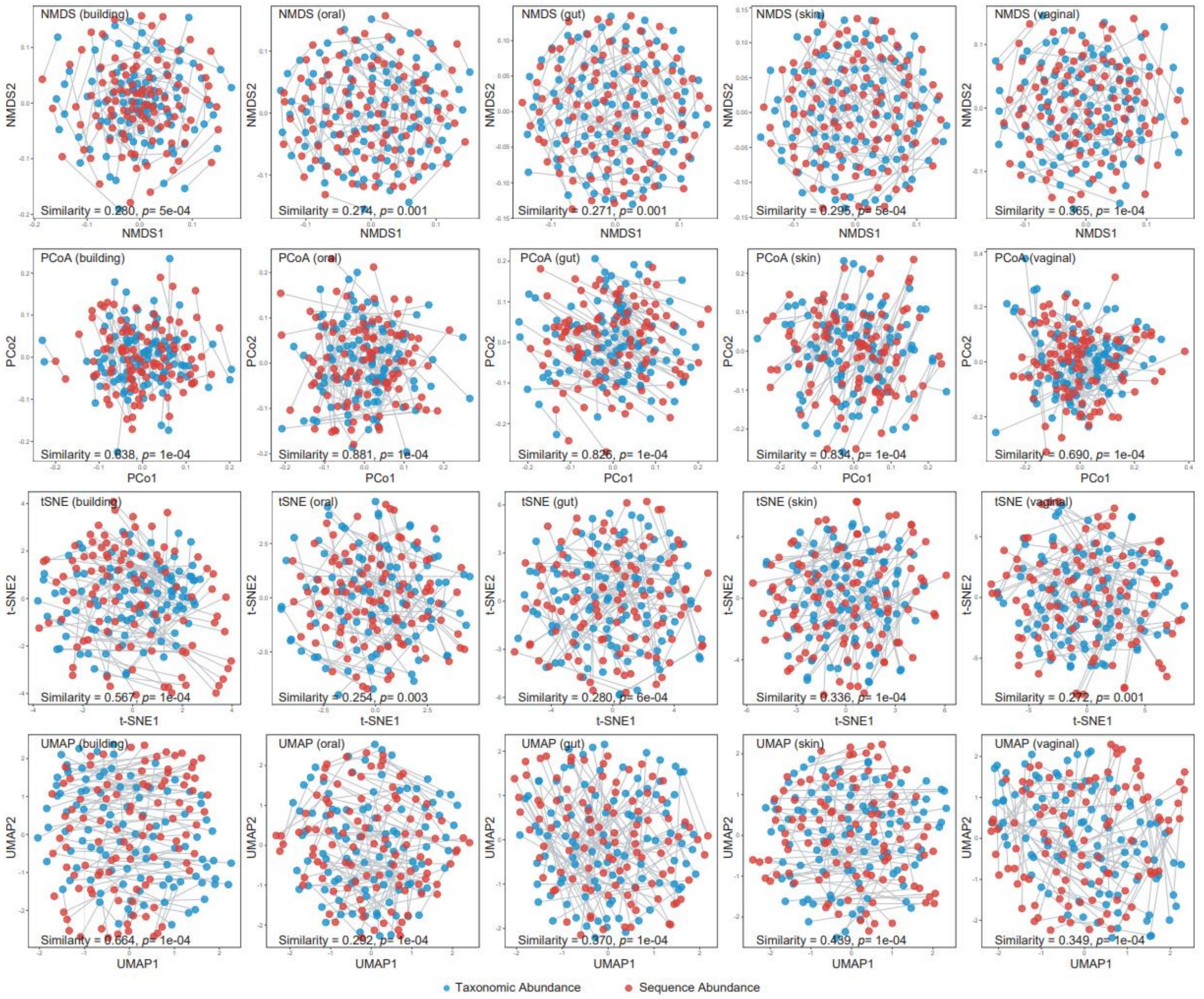


**Figure S2. Ordination analyses of simulated profiles based on BC.** Scatter plots of NMDS, PCoA, t-SNE and UMAP illustrate the dissimilarities between the sequence abundance (red dots) and taxonomic abundance (blue dots), which are the ground truth of the simulated profiles of 100 build environment, 100 oral, 100 skin, and 100 vaginal samples. Bray-Curtis (BC) distance was used to for the ordination analyses. The two abundance types from the same profile were connected using grey lines to show the difference of beta diversity.

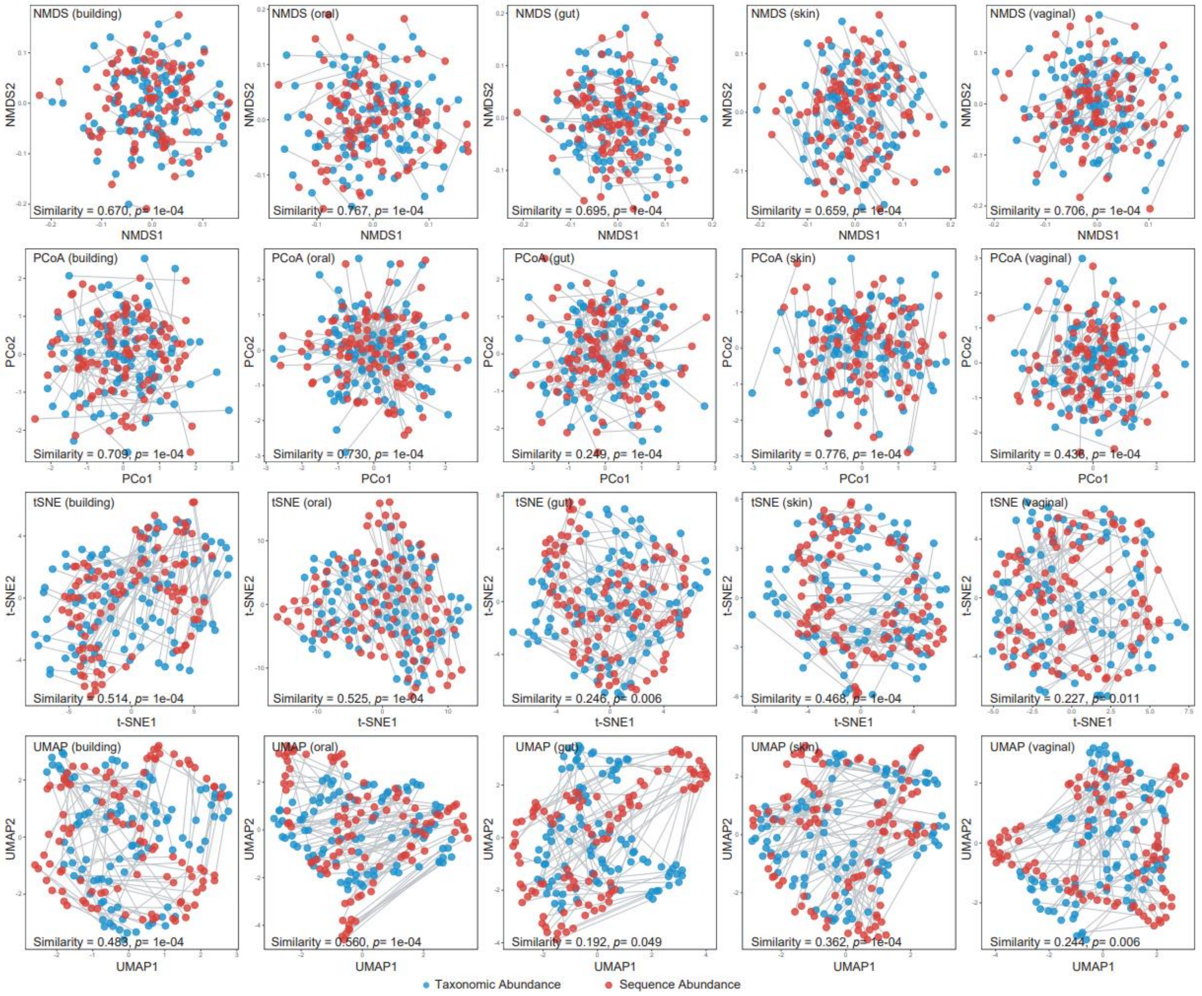


**Figure S3. Ordination analyses of simulated profiles based on rAD.** Scatter plots of NMDS, PCoA, t-SNE and UMAP illustrate the dissimilarities between the sequence abundance (red dots) and taxonomic abundance (blue dots), which are the ground truth of the simulated profiles of 100 build environment, 100 oral, 100 skin, and 100 vaginal samples. Robust Aitchison distance (rAD) was used to for the ordination analyses. The two abundance types from the same profile were connected using grey lines to show the difference of beta diversity.

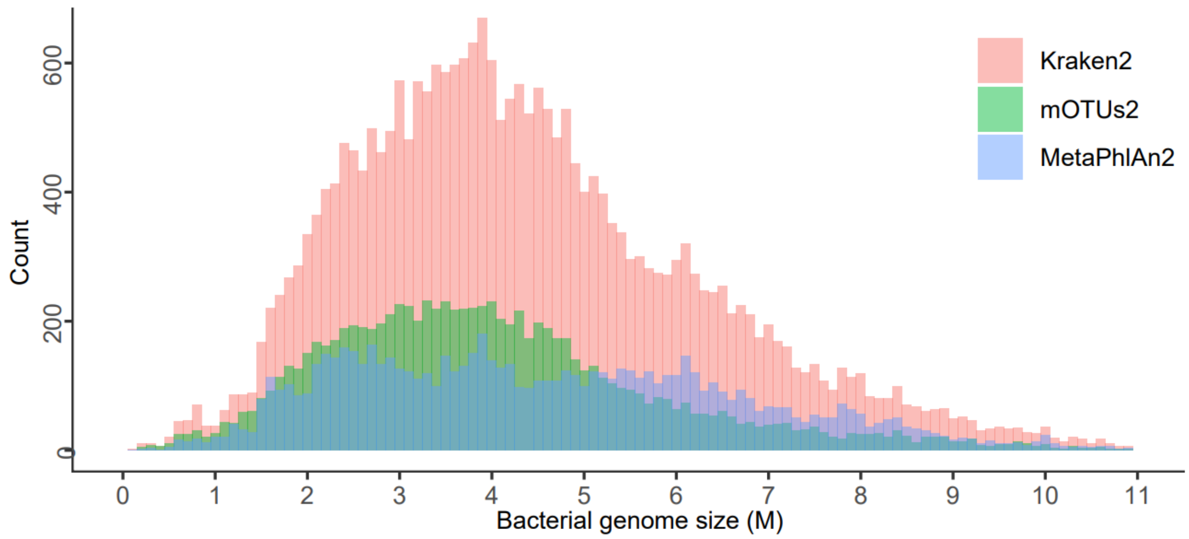


**Figure S4. Genome size distribution of bacteria in the reference databases used in Kraken2, MetaPhlAn2, and mOTUs2.**
